# Transcriptomic Analysis of the Spatiotemporal Axis of Oogenesis and Fertilization in *C. elegans*

**DOI:** 10.1101/2024.06.03.597235

**Authors:** Yangqi Su, Jonathan Shea, Darla DeStephanis, Zhengchang Su

## Abstract

The oocyte germline of the *C. elegans* hermaphrodite presents a unique model to study the formation of oocytes. However, the size of the model animal and difficulties in retrieval of specific stages of the germline have obviated closer systematic studies of this process throughout the years. Here, we present a transcriptomic level analysis into the oogenesis of *C. elegans* hermaphrodites. We dissected a hermaphrodite gonad into seven sections corresponding to the mitotic distal region, the pachytene, the diplotene, the early diakinesis region and the 3 most proximal oocytes, and deeply sequenced the transcriptome of each of them along with that of the fertilized egg using a single-cell RNA-seq protocol. We identified specific gene expression events as well as gene splicing events in finer detail along the oocyte germline and provided novel insights into underlying mechanisms of the oogenesis process. Furthermore, through careful review of relevant research literature coupled with patterns observed in our analysis, we attempt to delineate transcripts that may serve functions in the interaction between the germline and cells of the somatic gonad. These results expand our knowledge of the transcriptomic space of the *C. elegans* germline and lay a foundation on which future studies of the germline can be based upon.

## Introduction

With a transparent body of less than 1,000 somatic cells, a fully sequenced genome harboring 19,985 protein-coding genes (WS291 annotation) and about 14 hours of embryogenesis time and two weeks of life span, the *C. elegans* hermaphrodite provides an extraordinary model to understand various types of cell differentiation and organogenesis (Sulston and Horvitz, 1977, Sulston et al., 1983, Consortium, 1998, Hillier et al., 2005, Wood and Edgar, 1994, Rose and Kemphues, 1998, Labouesse and Mango, 1999, Kim et al., 2013, Chu and Shakes, 2013, Marcello et al., 2013, Robertson and Lin, 2013). Particularly, *C. elegans* gonad provides an excellent model to understand meiosis(Pazdernik and Schedl, 2013), gamete formation (Kim et al., 2013, Chu and Shakes, 2013) and fertilization (Marcello et al., 2013).

In the *C. elegans* hermaphrodite germline, oogenesis occurs independently in two sets of U-shaped gonads connected to a single shared uterus(Pazdernik and Schedl, 2013). Oocyte formation begins at the distal end of each gonad with mitotically proliferating germline stem cells near the single somatic distal tip cell (DTC). Proliferating germ cells away from the DTC begin to enter meiosis prophase I through a transition zone, after which germ cells move along the gonad while going through the pachytene, diplotene and diakinesis stages ending in the most proximal (-1) oocytes that awaits fertilization in the spermathecae for progression into metaphase I and the subsequent formation of the zygote. Apart from the proximal oocytes in diakinesis, most of the germline nuclei do not have fully enclosed membranes and form a syncytium, sharing a nucleus free cytoplasmic region called the rachis, which facilitates the transport of RNAs and proteins to growing oocytes (Figure 1A). Throughout this process, the germline also is enveloped by five pairs of gonadal sheath cells (Sh1-Sh5 from distal to proximal), each pair serving distinct functions through communication with the germline and promoting the oogenesis program(Pazdernik and Schedl, 2013).

**Figure 1.**
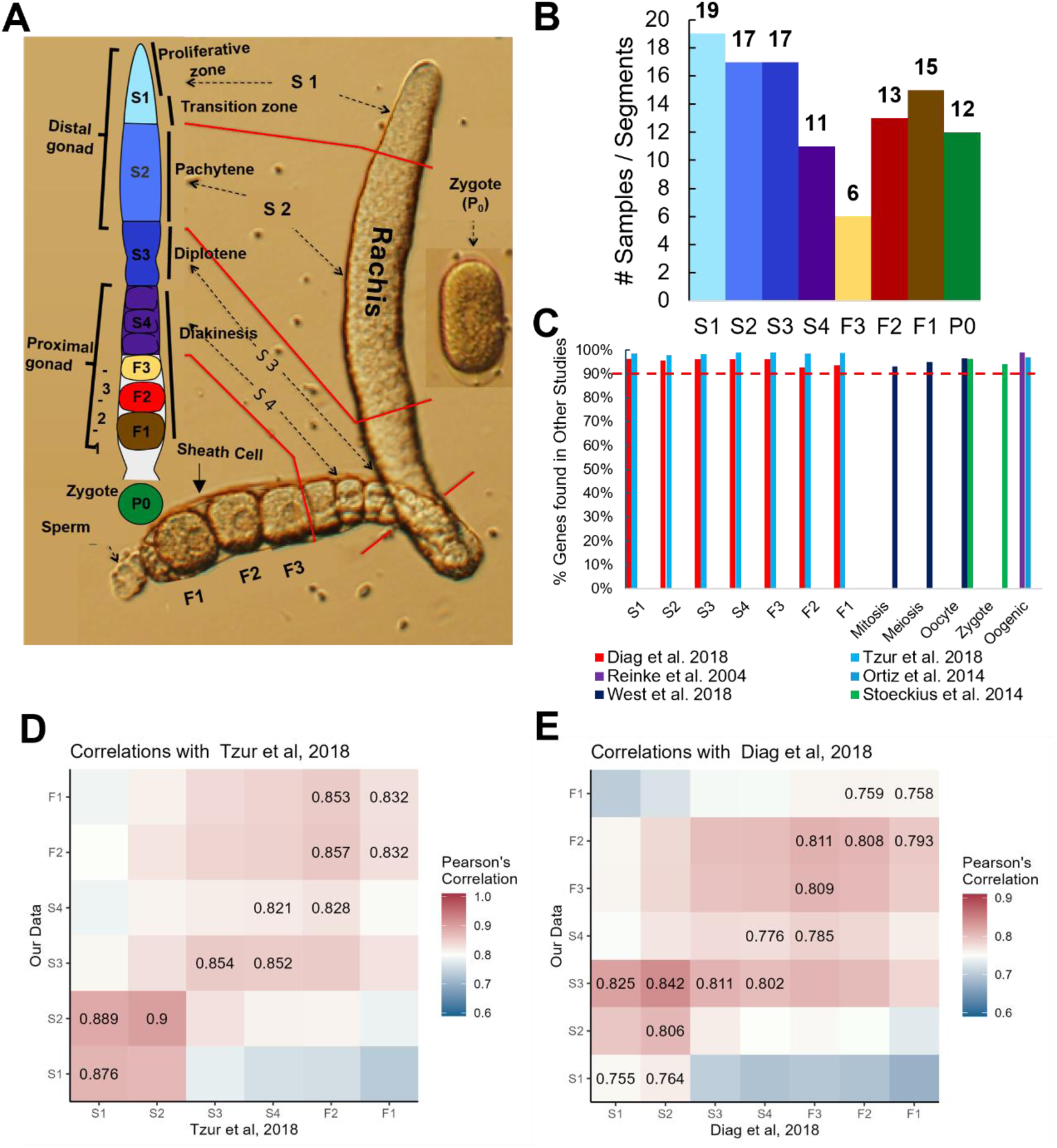
Comparison of our datasets with existing ones. A. A diagram of an isolated one side gonad together with a cartoon of one side gonad showing the dissection positions for the segments along the one side gonad. B. Number of samples from each stage of segments, oocytes and zygotes. C. Percentage of genes found expressed in each stage in previous studies that we found expressed in our study. D, E. Heatmap of Pearson correlation coefficient of our detected expressed genes in the segments with those of (Tzur et al., 2018) (D) and of (Diag et al., 2018) (E).

However, for a long time this system is limited by its miniscule size, preventing a detailed dissection of the biochemistry in each part of the oocyte assembly line using techniques such as transcriptome profiling using microarray (Reinke, 2002, Walhout et al., 2002, Baugh et al., 2003) or bulk-RNA sequencing (RNA-seq)(Spencer et al., 2011, Gerstein et al., 2010, Li et al., 2014), and proteome profiling using mass spectrometry (Yuet et al., 2015), as all of these techniques require a descent quantity of RNA/protein from at least hundreds of thousand cells.

Recent studies have performed micro-dissections of the *C. elegans* gonad and profiled transcriptomes of the gonad segments using single-cell RNA-seq (scRNA-seq) techniques (Diag et al., 2018). However, these analyses mainly focused on the post-transcriptional/translational regulation of germline transcripts via binding of 3’UTRs to RNA binding proteins and miRNAs. Although these studies provided expression estimates for genes from each segment as well, they did not focus on other aspects of transcriptomic changes between the segments that might also account for the progress of oogenesis. Consequently, we still lack a good understanding of the machinery of the assembly line, such as key regulators and gene expression patterns along the temporal and spatial axis of the gonad.

To fill these gaps, we combined microdissection with scRNA-seq technique(Tang et al., 2010a, Tang et al., 2010b, Ramskold et al., 2012, Picelli et al., 2013), and profiled the transcriptomes in the proliferative zone, pachytene zone, diplotene zone, early diakinesis zone (before -3 oocyte stage), later diakinesis zone (-3, -2, -1 oocytes), and the zygote. Our results revealed a highly dynamic picture of gene transcriptional regulation at each transitional time point throughout the oocyte assembly line. These results should provide a foundation to further understanding the molecular mechanisms of the oogenesis and fertilization processes.

## Results

### Expression levels of detected genes correlate well with those from previous studies

We cut each isolated gonad into seven segments roughly corresponding to the stages of oocyte development (Figure 1A) (Materials and Methods), and the number of samples collected for each segment, oocyte and the zygote are shown in Figure 1B. To assess the quality of our RNA-seq libraries, we evaluated the similarity between the detected genes and their expression values and those from six previous studies (Reinke et al., 2004, Ortiz et al., 2014, Stoeckius et al., 2014, West et al., 2018, Tzur et al., 2018, Diag et al., 2018) (Materials and Methods). Four (Reinke et al., 2004, Ortiz et al., 2014, Stoeckius et al., 2014, West et al., 2018) of these studies largely quantified expression levels in entire gonads or large sections of the gonad, thus we aggregated gene expression in corresponding samples to allow reasonable comparisons. Our aggregated expression profiles recall over 90% of expressed genes in all the four datasets (Reinke et al., 99%; Ortiz et al., 97%; West et al., 93% mitotic, 95% meiotic and 96% proximal oocyte; Stoeckius et al., 96% oocyte and 94% zygote) (Figure 1C).

Two of these studies (Tzur et al., 2018, Diag et al., 2018) dissected the *C. elegans* gonad into multiple segments and profiled the transcriptome of each segments using a variety of techniques including RNA-seq. As both studies cut the gonad in more segments than we did, we aggregated data from the segments of (Tzur et al., 2018) and (Diag et al., 2018) according to the alignments of the segments (Materials and Methods, Supplementary Tables S1), so that data from largely the same segments as ours were compared. Our detected genes in each segments and oocytes recall most of detected genes in the corresponding aggregated segments by (Tzur et al., 2018) and (Diag et al., 2018) (Figure 2C). Moreover, the expression levels of genes in our segments are largely correlated with those in the corresponding aggregated segments in the two prior studies (Figures 1D, 1E). These results suggest that we have largely correctly align the segments in both studies to ours. However, notably, our detected genes have higher recall rates for (Figure 1C) and higher correlation coefficients with (Figures 1D, 1E) those of (Tzur et al., 2018) than for and with those of (Diag et al., 2018). This might be due to the more similarity between our segments and those of (Tzur et al., 2018) than between our segments and those of (Diag et al., 2018). Taken together, these results suggest that our detected genes are largely consistent with those detected by previous studies.

**Figure 2.**
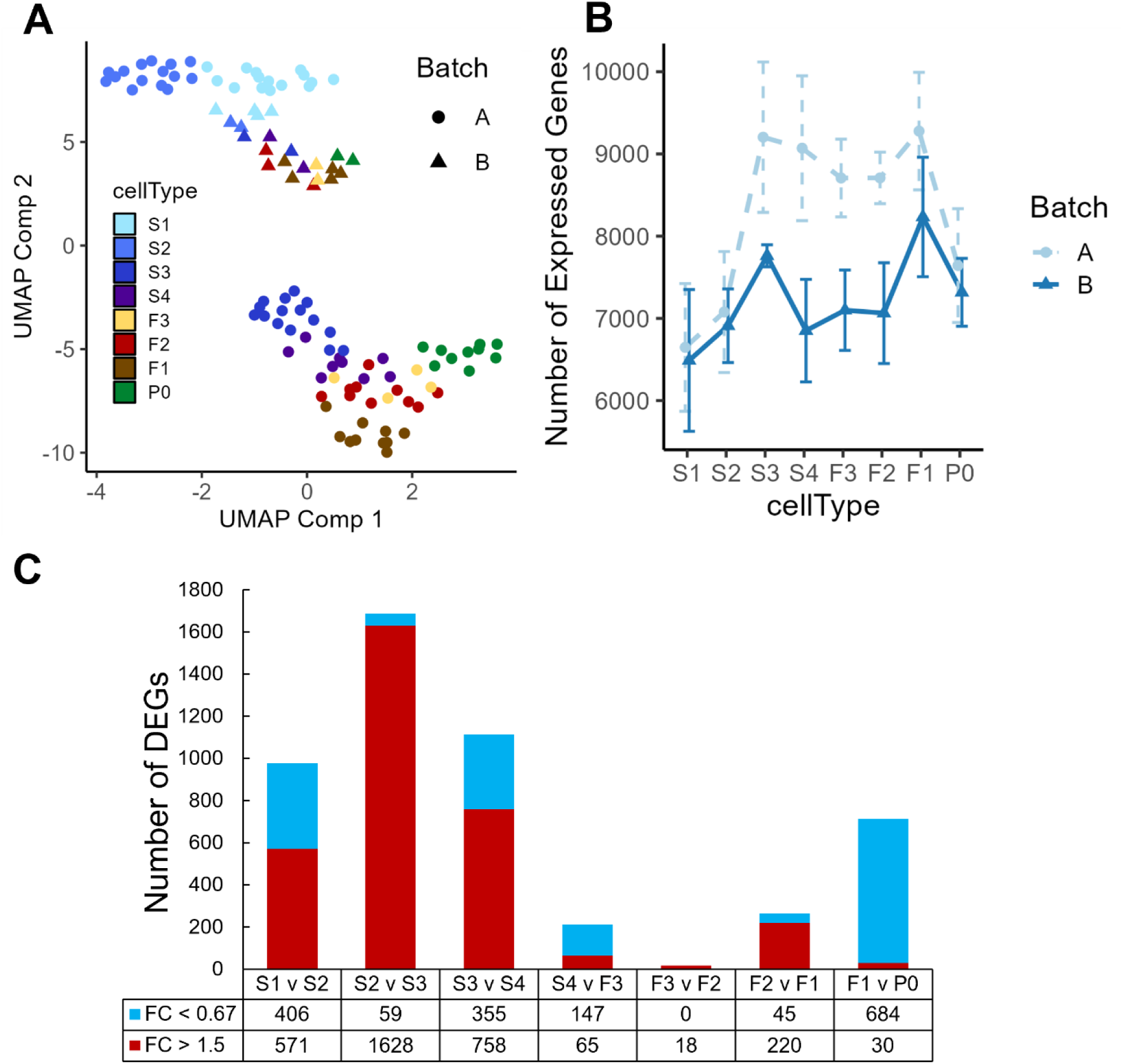
UMAP display of samples and differential expression analysis of genes between neighboring stages. A. Both batches of samples are clustered according to their positions along the gonad developmental axis by UMAP based on their measured transcriptomes. B. Boxplot of numbers of genes detected in the samples in each developmental stage of the gonad and zygotes. C. Number of upregulated and downregulated genes detected between each pair of neighboring stages, see Supplementary Table S3 for details.

### Differential gene expression occurs in early stages of oogenesis and mostly in proximal oocytes

Dissection of the *C. elegans* hermaphrodite gonad is a delicate procedure that is prone to contamination from neighboring tissues due to the miniscule size of the gonad and proximity of neighboring cells such as sheath cells, intestine cell and released sperms. To mitigate the effects of such contaminations, we filtered sperm-, intestine- and stress-related genes as well as heavily contaminated samples (Materials and Methods, Supplementary Figure S1, Supplementary Table S2). Mover, the hermaphrodite gonad itself also contains somatic cells, most notably the five pairs of gonadal sheath cells that tightly enclose the germline. The sheath cells mainly function to provide germline maturation signals, move germ cells along the rachis and push proximal oocytes into the spermathecae. Due to the tight interactions between the sheath cells and the germline, complete removal of these cells was very difficult, especially for earlier stages (S1-S4). Thus, some differential gene expression results for these early stages are inevitably due to differences between gonadal sheath cells, albeit they seem to negligibly affect our results for these stages of comparison as described below.

We inspected the relationships among our samples via UMAP visualizations. As shown in (Figure 2A), the samples form into two distinct clusters, indicating strong batch effects in our datasets possibly due to the two different scRNA-seq library preparation protocols used at different stages of the project (Materials and Methods). Nonetheless, a trajectory from S1 samples to F1 and zygote samples is formed in both batches, which is in line with the developmental path of the germline. Thus, we account for batch effects in subsequent analysis when possible. Inspection of the number of genes expressed in each segment/cell type shows a clear pattern, i.e., the number of expressed genes increased from S1 to S3, before dropping slightly in S4 and exhibiting only minor changes before another increase in the -1 oocyte (F1) and finally a large decrease in the fertilized oocyte (Figure 2B). Therefore, it appears that gene transcriptional regulation mostly occurs in early stages of oogenesis, particularly between the S2 (pachytene) and S3 (diplotene) transition and becomes progressively quieter as the oocyte goes through the S4, F3 and the F2 stages (Figure 2B). Gene transcription are reactivated in the F1 oocytes, probably preparing for fertilization (Figure 2B). To further reveal gene expression transitions alone the developmental axis of the gonad, we analyzed DEGs between each pair of neighboring stages with the earlier stage as the baseline reference in each comparison (Figure 2C). Transition from S2 to S3 invokes the largest number of up-regulated DEGs, and transition from F3 to F2 has the smallest number of DEGs, while fertilization triggers the largest number of downregulated DEGs in the zygotes (Figure 2C).

### DEGs form distinct clusters that are significantly enriched for various functions related to oogenesis

To reveal functional modules underlying the maturation process and fertilization of oocytes, we clustered the union of DEGs identified in all neighboring stages comparisons, based on their expression levels in all analyzed samples. As shown in Figure 3, the DEGs form distinct clusters that are significantly enriched for various functional modules. For instance, clusters 2, 4 and 6 are significantly enriched for ribosomal and translation related processes. All these three clusters of genes exhibited a downregulating trend of expression, albeit with their largest decrease at different stages. Cluster 14 and 15 are enriched for genes involved in programmed cell death, with expression levels elevated in in the S3 stage corresponding to the diplotene loop. However, genes in cluster 14 were quickly downregulated after the S4 stage, while genes in cluster 15 retained similar transcription levels through the subsequent stages. Cluster 18 -20 are all enriched for processes related to oogenesis, e.g., eggshell formation and female gamete generation. Genes in these three clusters exhibited increasing trends of expression from S1 to -1 Proximal oocyte (F1), with the largest increases happening in the early stages (S1-S3). However, genes in cluster 18 experienced reduced expression after fertilization in the zygotes (P0 cells), while genes in cluster 19 and 20 remained similar expression levels. Furthermore, genes in cluster 18 are enriched for eggshell formation, suggesting that transcripts-related to eggshell formation begin degradation post-fertilization after their protein products are no longer needed. Most DEGs belonging to the larger clusters 16 and 17 exhibited similar increases in expression from S2 to S3 and maintained steady levels of expression throughout the later stages even post fertilization. These genes are involved in phosphorylation, synaptic transmission and signaling, positive regulation of transcription, neuronal differentiation, cell fate specification and cell migration. Gene involved in cell migration might be responsible for the mobility of oocytes along the rachis. Interestingly, cluster 17 is strongly enriched for genes involved in neuronal development, suggesting common functional modules might be used in the differentiation processes of both neurons and oocytes. Cluster 9 is enriched for genes involved in muscle structures and myofibril assembly. As mentioned above, proximal gonadal sheath cells serve the role of pushing oocytes into the spermathecae and require many components like those of muscle cells. Thus, it is highly likely that genes of this cluster originate from contamination of proximal sheath cells wrapped around the proximal oocytes. It is also worth noting that gene expression pattern of cluster 9 differ from those of clusters 16 and 17 in that expression of genes in cluster 9 almost completely disappears in fertilized zygotes, likely due to the absence of sheath cells surrounding the isolated zygotes. Cluster 1 exhibits no obvious pattern of change in expression and the expression levels are generally low. These genes are enriched for defense response related processes and might be required at low levels along the gonad temporospatial axis. Both clusters 5 and 11 are enriched for extracellular matrix organization. It has been shown that many genes (*mig-6, mig-39, lag-2, let-2, epi-1*, etc.) in the two clusters (Supplementary Table S4) were preferentially expressed in the distal mitotic regions of the gonad and played roles in extracellular matrix organization and distal tip cell migration (Henderson et al., 1994, Huang et al., 2003, Kawano et al., 2009, Kikuchi et al., 2015). Consistently, expression levels of these genes were elevated in S1.

**Figure 3.**
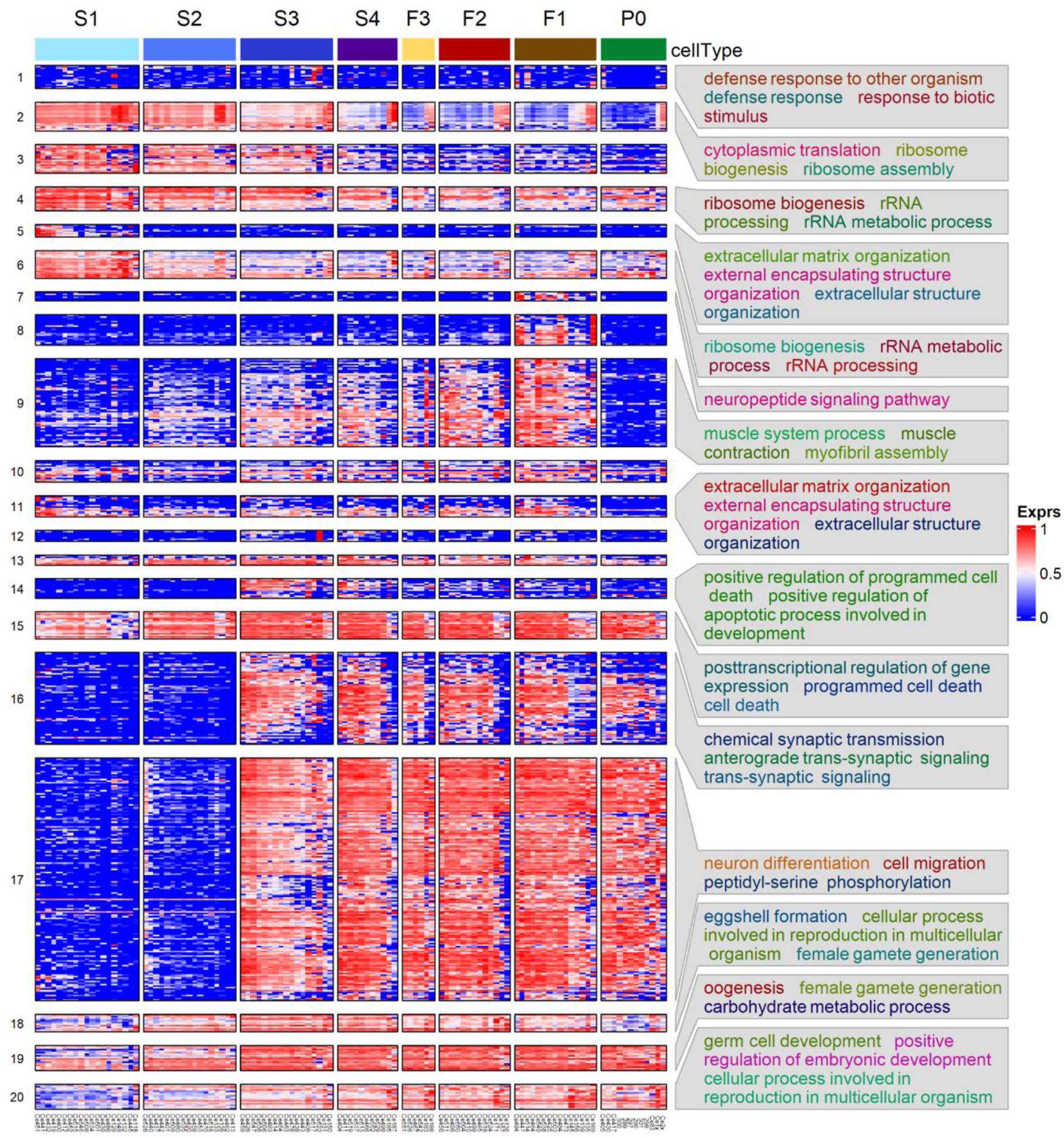
Heatmap of hierarchical clustering of DEGs using their transcription levels across the seven stages of oogenesis and zygotes. Enriched GO biological processes in some clusters are shown. See Supplementary Table S4 for details.

We also performed Gene Set Enrichment Analyses (GSEA) using shrunken Log2FC values of all genes evaluated between each pair of neighboring stages and the results are summarized in Supplementary Table S5. Although most of the GSEA results are in accordance with those observed in the gene clustering enrichments (Figure 3), surprisingly, GSEA finds upregulated genes enriched for cell cycle activity, mitosis, transcription, mRNA splicing, mitochondrial translation, and ATP production in the F1 vs P0 comparison (Supplementary Table S5). This suggests that transcriptional activation of cell division and energy production is present in the zygote.

### Possible contaminations of sheath cells in proximal oocytes samples

As mentioned above, there are possible contaminations in the proximal oocytes (F1∼F3) samples from surrounding somatic cells. This presents a challenge in deciphering whether expression changes originate from the germline or from the surrounding somatic tissues. As zygotes were often released in the medium once a cut was made across the vulva, and were always collected without obvious objects wrapped around, thus the zygote sample were unlikely contaminated by surrounding somatic cells. Therefore, we postulate that genes that are detected in proximal oocytes (F1∼F3) samples but absent in zygote samples (such as those found in clusters 7-9 in Figure 3) are likely from contaminating tissues, and find many gene meet this criterion. Most notably, expression of *let-23* and *itr-1* were relatively stable between F2 and F1 prior to dropping significantly in the zygotes, while expression of lin-3 remained high and relatively unchanged between proximal oocytes and zygotes (Figure 4A). The contractile activity of sheath cells begins with major sperm protein signals to the proximal oocytes, which in turn produces and releases LIN-3 ligands that are received by the LET-23 receptor on proximal sheath cells (Miller et al., 2001). The LET-23 receptor then triggers signaling inside the sheath cells through PLC-3, which phosphorolyzes IP3 that binds to ITR-1 receptors on the ER, causing the release of calcium(Yin et al., 2004). In addition, sheath cell specific innexin channel encoding genes *inx*-8 and *inx*-9 (Starich et al., 2014) maintained intermediate expression levels in S1∼S4 stages, and were highly upregulated in proximal oocytes, but had negligible expression levels in the zygotes (Figure 4BA). Furthermore, expression levels of sheath cell contractile activity related genes were also progressively increased along the gonadal development axis, but almost vanished in zygote samples, such as genes *pat*-10, *mup*-2, *tni*-1 and *unc*-27 coding for the troponin complex (Obinata et al., 2010, Ono and Ono, 2004)(Supplementary Figure 2A), and genes *unc*-54 and *myo*-3 coding for the myosin heavy chain (Ono and Ono, 2016, Shelton et al., 1999) (Supplementary Figure 2B).

**Figure 4.**
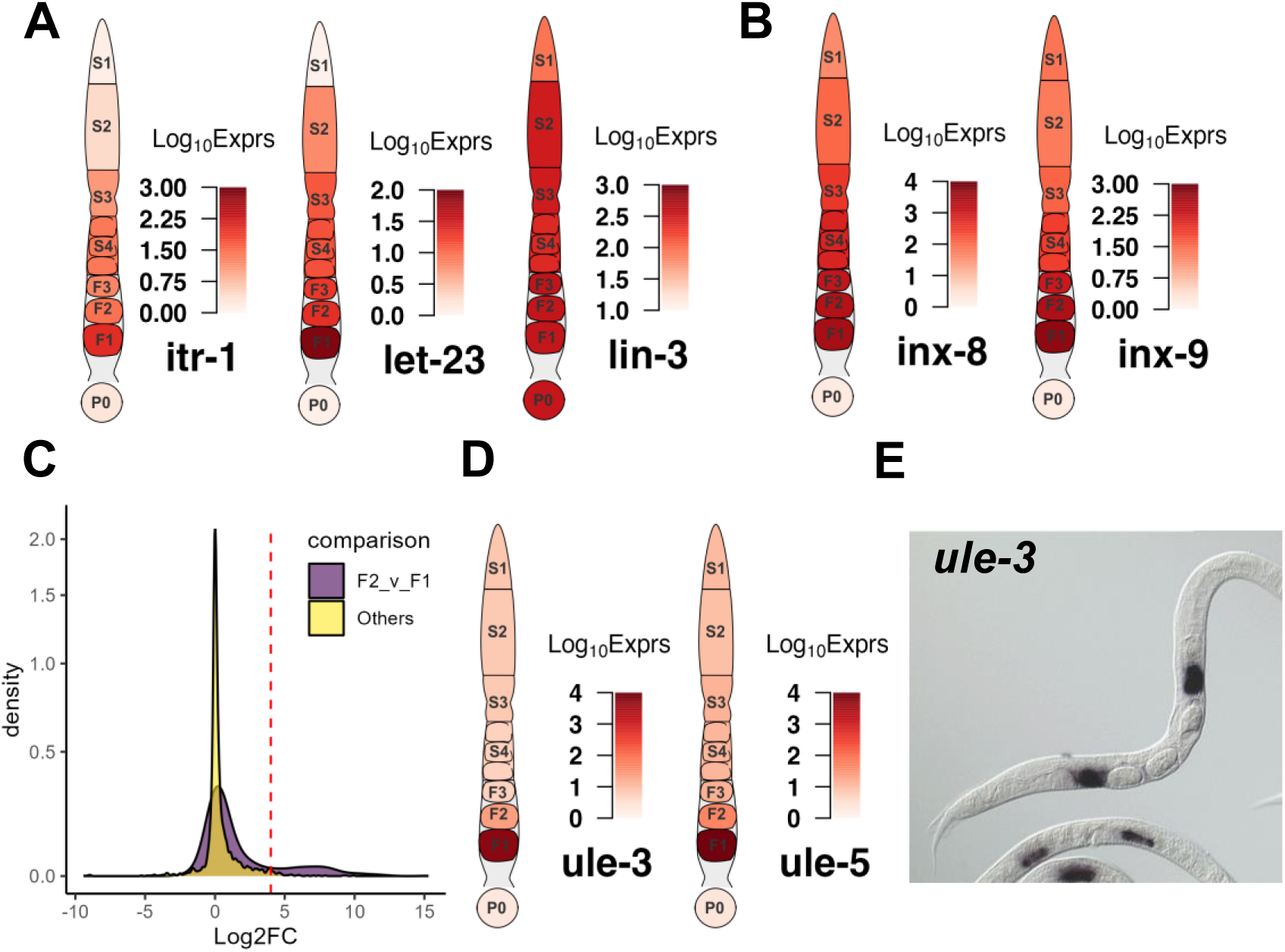
Examples of transcriptional dynamics of possible sheath cell genes along the gonad developmental axis. A. Genes coding for hemichannels (*inx*-8 and *inx*-9) of the somatic gonad. B. Genes coding for components signaling pathways between proximal sheath cells and oocytes. C. Distribution of Log_2_FC values between neighboring stages of the DEGs that are significantly downregulated in the F1 vs P0 comparison. A small portion of these DEGs is significantly upregulated in the F2 vs F1 comparison as indicated by the right peak of the distribution compared to other comparisons, see Supplementary Table S6 for details. D. Genes coding for ULE-3/5. E. NEXTDB in situ imaging of ule-3 expression in spermathecae. In each gonad diagram, the average expression levels of the genes in each segment or the zygote are shown.

Similar reasoning can be made with other genes that have previous evidence of somatic or germline origins. Searching the CenGEN database(Hammarlund et al., 2018) revealed that genes *perm-2/4* encoding components of the eggshell(Gonzalez et al., 2018) had the highest expression levels in sheath cells. Expression of perm-2/4 is absent in P0 but high in F1 (Supplementary Figure 2C), while other known components of eggshell that are produced in the germline such as egg-1/2 do not exhibit such significant decreased expression in P0 (Kadandale et al., 2005). The expression of myosin light chains genes *mlc-1/2/3* involved muscle activity (Rushforth et al., 1998, Moerman et al., 1997) are all high in F1 but absent in P0, while the expression of the non-muscle myosin light chain gene *mlc-4* required for cytokinesis in zygotes, is present in P0 (Supplementary Figure 2D) (Ono and Ono, 2016, Shelton et al., 1999). Analysis of actin genes *act-1/2/3/4* (Ono, 2014, Ono and Pruyne, 2012) may even suggest that the expression of act-4 is not required in zygotes, as it is the only actin gene with negligible expression in P0 (Supplementary Figure 2E), an observation also supported by previous findings that *act-1/2/3* were expressed in both muscle and non-muscle cells, while *act-4* was expressed predominantly in body wall muscle(Stone and Shaw, 1993, Willis et al., 2006).

### Proximal oocyte expression profiles reveal interactions between the germline and the somatic gonad

We compared the distributions of Log2FC values for all the comparisons of the DEGs that exhibit significantly lower expression levels in P0 samples in the F1 vs P0 comparison. As shown in Figure 4C, a considerable number of the genes show a significant increase in expression between the F2 vs F1 comparison, as indicated by an additional small peak with higher Log_2_FC values in the distribution compared to other comparisons. These genes might account for those in clusters 7 and 8 (Figure 3). This is interesting, as early studies indicate transcriptional inactivity or an overall presence of transcriptional silence as oocytes move to the proximal end (Walker et al., 2007, Starck, 1977). Though this was the case for the F3 to F2 transition, however, clearly not for the F2 to F1 transition (Figure 2C). Two Uterine Lumen-Expressed (ule) genes *ule*-3 and *ule*-5 (Figure 5D) exhibited sudden increases in transcription from 10-fold to 100-fold between the F2 and F1 transition. This is different from expression patterns of genes of gonadal sheath origin that we described earlier, where the expression levels stay relatively stable in the proximal oocytes. It has been reported that *ule*-3/5 might play a role in driving the ageing of the reproductive system, though the origin of their expression is not clearly discernable (Zimmerman et al., 2015). A more recent study utilizing FISH to track the origins of these transcripts suggests a mechanism by which the transcripts are produced in spermathecae and carried over into the proximal oocytes(Trimmer et al., 2023). Using the expression of ule-3/5 as a reference, we discerned a set of 25 genes displaying the similar expression pattern (Supplementary Table S6) by invoking an stringent criteria: Log_2_FC < -7 in the F1 vs P0 comparison; and Log_2_FC > 4 in the F2 vs F1 comparison; and average Median Normalized Expression in F1 > 1000). From the NEXTDB database(Kohara, 2001), we were able to obtain in situ hybridization imaging for the products of 17 of these genes, of which 13 genes exhibited clear localization of corresponding transcripts in the spermathecae region (Figure 4E, Supplementary Figure S3). These results suggest possible interactions between transcriptionally silent oocytes and its somatic neighbors, where the transcriptional events happen in the surrounding spermathecae, and the proximal oocytes take up the transcripts and produce protein products.

**Figure 5.**
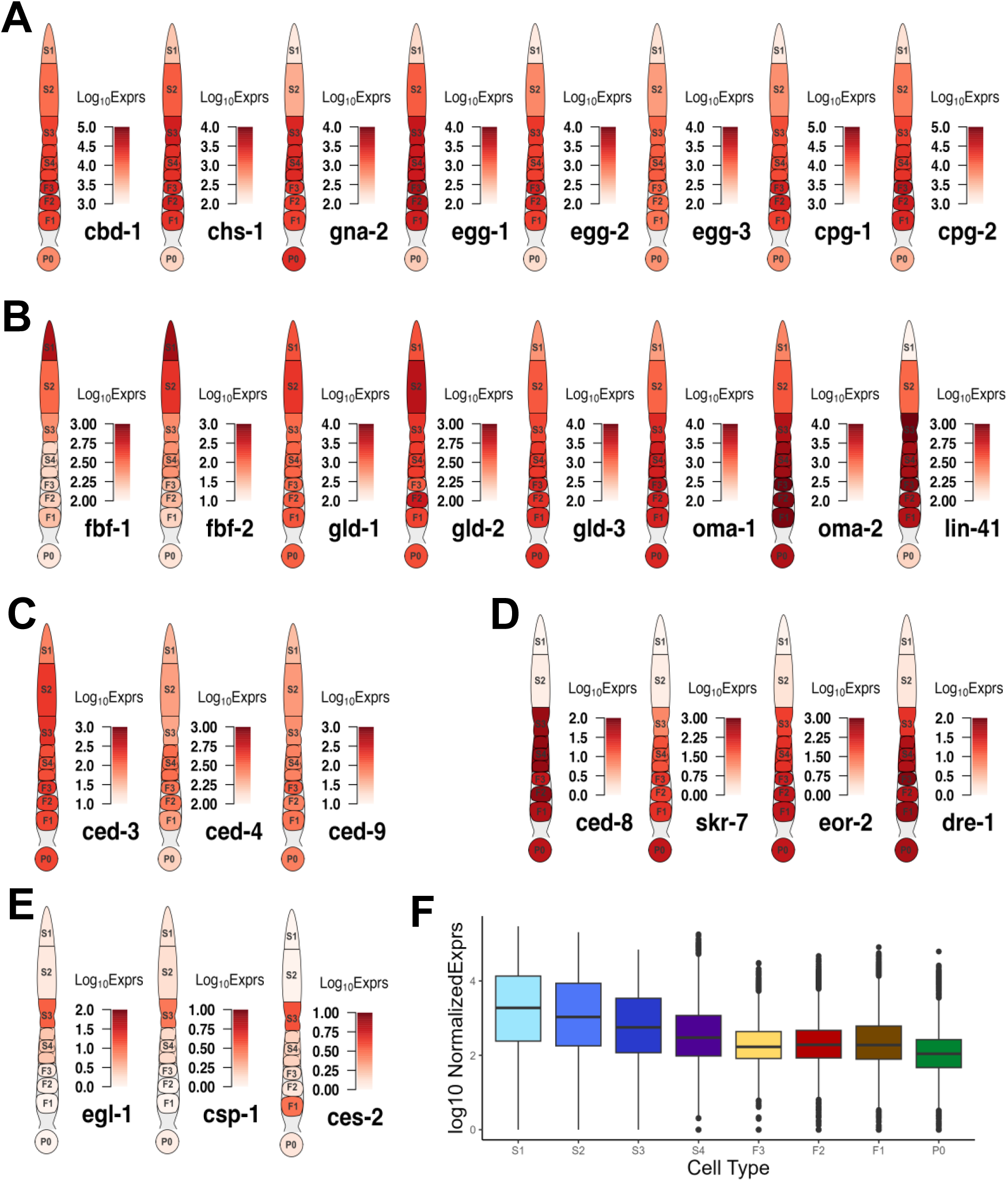
Examples of transcription of DEGs that are involved in key events of oogenesis and fertilization. A. Genes encoding different elements of the eggshell. B. Genes involved in mitosis-meiosis transition and meiotic maturation. C. Genes involved in apoptosis with expression throughout the gonad. D. Genes involved in apoptosis with elevated expression starting from the S3 stage. E. Genes involved in apoptosis showing transient expression in the S3 stage. F. Boxplot of transcription levels of genes coding for ribosomal subunit across each stage of gonad development and in zygotes.

### DEGs mark transcriptional timing of the key events of oogenesis and fertilization

One of the early key events in the oogenesis process is the control of mitosis and meiosis. Thus, it is interesting to look into the transcription patterns of mitosis promoting regulators FBF-1/2 (Crittenden et al., 2002) and meiosis promoting regulators GLD-1/2/3 (Eckmann et al., 2004). We found *fbf-1/2* showed high expression in the S1 stage, but lower expression in the S2 and S3 stages and beyond (Figure 5A), while *gld*-2/3 exhibited increased expression only between S1 and S2, and then along with *gld*-1 maintained high expression in the later stages (Figure 5A). In addition, we observed transcriptional regulation of key factors involved in the maintenance of germ cells in meiotic prophase I such as OMA-1/2 and LIN-41(Tsukamoto et al., 2017). Specifically, the expression of *oma*-1/2 and lin-41 gradually increased throughout the early stages (S1 and S2) of oogenesis followed by high elevations in the S3 stage, which were maintained even after fertilization, apart from *lin*-41, whose expression dropped after fertilization (Figure 5A). It has been suggested that LIN-41 could prolong prophase I and inhibit meiotic maturation after fertilization by a translational level regulatory mechanism(Spike et al., 2014, Tsukamoto et al., 2017), thus diminishment of the *lin-41* transcription in zygotes suggests that transcriptional degradation might also play a role in the exit of the oocyte from metaphase I upon fertilization.

We also found that many genes coding for eggshell components were upregulated in distal segments of the gonad, far before the complete formation of the eggshell that happened around the early-stage embryo (Stein, 2018). Genes coding for components of the vitelline layer (cbd-1)(Gonzalez et al., 2018), the chitin layer (chs-1, gna-2, egg-1/2/3) (Zhang et al., 2005, Johnston and Dennis, 2012, Johnston et al., 2006, Kadandale et al., 2005, Maruyama et al., 2007) and the proteoglycan layer (cpg-1/2) (Olson et al., 2006) all exhibit increased expression in early stages of the germline until after fertilization (Figure 5B). These results suggest that transcription of these eggshell genes occur mostly during the mitosis to meiosis transition and the pachytene, while translation and degradation of these transcripts might occur as a response to fertilization signaling.

Moving along the germline, another key event of oogenesis happens in the diplotene loop (S3) where germ cells undergo apoptosis. Here we find that genes regulating apoptosis form three distinct patterns of expression. The expression of genes encoding core apoptosis machinery such as apoptosis initiators CED-4/3(Huang et al., 2013) and apoptosis inhibitor CED-9 (Hengartner et al., 1992) were initially low in the S1 stage but elevated to steady states in the S2 (pachytene) stage (Figure 5C). The high expression levels of both *ced*-3 and *ced-*9 were largely maintained thereafter until after fertilization, while that of *ced*-4 was maintained thereafter until the F2 stage and then gradually decreased in F1 and zygotes (Figure 5C).

Expression of *ced*-8, which encodes a substrate of the CED-3 Caspase and is likely involved in regulating the timing of apoptosis (Chen et al., 2013), follows a different pattern, with significant upregulation in the S3 stage, and maintaining high expression until fertilization (Figure 5D). The sudden increase in *ced*-8 transcription in the S3 stage suggest that CED-8 might play an important role in initiating apoptosis in the germline. Other genes such as *skr*-7, *eor*-2 and *dre*-1 showed expression patterns like that of *ced*-8, with elevated expression starting from the pachytene (S3) loop onwards through fertilization (Figure 5D). SKR-7 has been implicated in inducing apoptosis(Gao et al., 2008), and DRE-1 has been found to interact directly with CED-9 in regulating apoptosis(Chiorazzi et al., 2013). Early studies have found EOR-2, along with EOR-1 to induce apoptosis in neuronal cells(Hoeppner et al., 2004). However, we only observed upregulation of *eor-2* (Figure 5D) but not of *eor-1* in the germline, suggesting a different mechanism of EOR-2 induced apoptosis in the germline than in neuronal cells.

The third group of apoptosis related genes follow a different expression pattern that can be characterized by the expression profile of *egl-1*, which encodes a direct downstream target of CED-4 and inhibitor of CED-9, playing a critical role in DNA damage induced germline apoptosis (Huang et al., 2013). *Egl-1* exhibited a transient increase in transcription in the pachytene loop (S3) that did not go beyond the S4 stage (Figure 5E). Other apoptosis related genes such as *csp*-1 and *ces*-2 displayed expression patterns like that of *egl*-1 (Figure 5E). An earlier study has found that *csp-1* was expressed in late stage pachytene of the germline using FISH imaging (Denning et al., 2013). *Ces-2* has been implicated in the apoptosis of neuronal cells in *C. elegans*, though a previous study suggested that *ces*-2 was not essential for germline apoptosis (Metzstein et al., 1996). However, the sudden upregulation of *ces-2* transcription in S3 strongly suggests a role of *ces-2* in apoptosis of the germline. Furthermore, since all 3 genes belong to cluster 14 (Figure 3), it is likely cluster 14 contains other genes that are related to apoptosis as well.

As shown in Figure 5F, genes encoding ribosome subunits and other translation-related proteins generally exhibited downtrends in transcription as oocytes matured and prepared for fertilization, consistent with a previous observation (Diag et al., 2018). Interestingly, genes downregulated between S1 and S2 are mainly involved in ribosomal precursor production, such as *eif-6, rpoa-1/2, fib-1, nucl-1*, etc. (Supplementary Table S3) (Miluzio et al., 2009, Xu et al., 2023), while those downregulated between S4 and F3 mainly encode ribosomal protein subunits, such as *rla-0/1, rpl-1/2/3/4/5/7/9/10/13/14/15/16/17* and *rps-0/1/2/3/4/5/7/8/9/10/11/12/13/14/15* (Supplementary Table S3) (Nakao, 2004). This suggests that the preparation of ribosomal assembly machinery for oogenesis mainly occurs in the distal mitotic regions prior to entering pachytene (S2), but ribosomal proteins continue to be produced until diakinesis (S4). Moreover, the reduced levels of expression from the S4 stage and beyond indicating that all the transcription of translational machinery required for oocyte maturation are formed before the diakinesis stage.

### Differential Alternative Polyadenylation activity resumes post-fertilization

Though a great deal of literature has focused on regulation of translation through the 3’UTRs of transcripts by ribosomal binding proteins (RBPs), few have elucidated changes of the 3’UTRs themselves (Merritt et al., 2008, Mangone et al., 2010a, Diag et al., 2018, Steber et al., 2019). Thus, we analyzed differential alternative polyadenylation (DAP) usage through the DaPars2 software (Feng et al., 2018, Li et al., 2021), which estimates changes in proportion of distal (lengthened 3’UTR) and proximal (shortened 3’UTR) polyadenylation sites used in two conditions. We found very few significant changes in distal versus proximal sites usage between neighboring stages, apart from the S4 vs F3, F3 vs F2 and F1 vs P0 comparisons (Figure 6A). GO term enrichment analysis found that only the F1 vs P0 comparison resulted in significant enrichment of genes with ADP for mitotic cell cycle related processes, with mostly shortened 3’UTRs (Figure 6B). For instance, we find that *cyb*-1/2.2 exhibit shortened 3’UTRs while *cdk*-1 exhibited lengthened 3’UTR (Figure 6C). CYB-1/2.2 along with CDK-1 regulate M phase entry of cell cycle in *C. elegans* (Rabilotta et al., 2015). Though most DAP genes between F1 and P0 exhibit shortened 3’UTR (Figure 6B), it is not straightforward how usage of distal vs proximal sites functions in the regulation of the final expression of protein products. Furthermore, despite very few significant DAP genes in the early stages of oogenesis, we found par-5 gene to exhibit DAP in both the S1 vs S2 and F2 vs F1 comparisons (Figure 6C). In fact, 3’UTR length of the par-5 transcript gradually decreased until F3 stages before increasing again (Figure 6C). It has been reported that PAR-5 regulates asymmetric cell division and alternative 3’UTR isoforms of par-5 confers different levels of PAR-5 protein abundance (Mikl and Cowan, 2014). Interestingly, most significant DAP genes between S4 and F3 exhibited an increase in 3’UTR length, while most DAP genes between F3 and F2 exhibit decreased in 3’UTR length (Figure 6A). However, it is unclear whether this is because F3 oocytes are fully cellularized and maintain a stable transcriptome or other factors.

**Figure 6.**
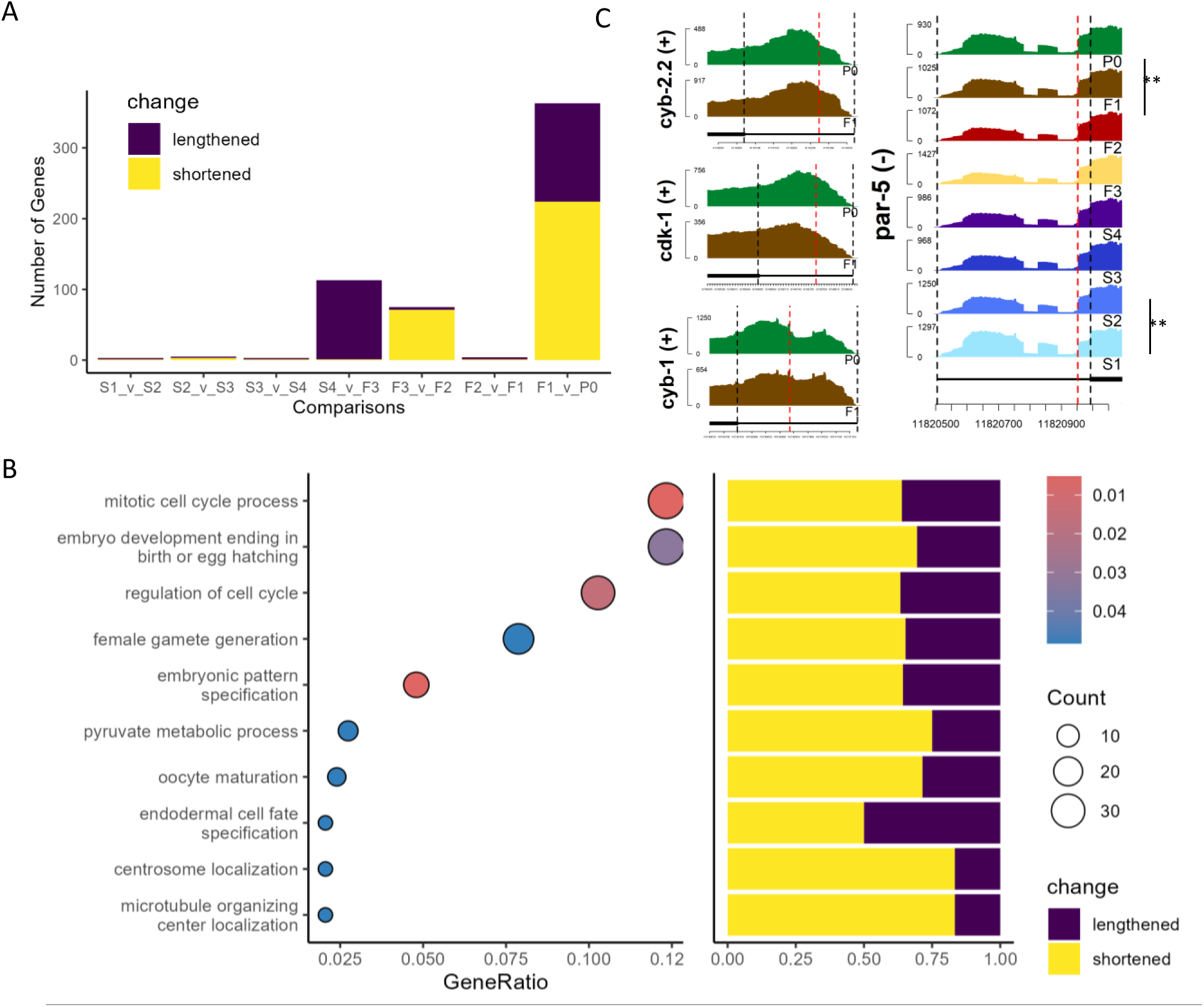
Differential alternative polyadenylation (DAP) analysis of genes between neighboring stages. A. Number of DAP genes between each neighboring stage, colors indicate lengthening (purple) or shortening (yellow) of 3’UTR lengths. B. GO term enrichment of significant DAP gene between F1 and P0 (left panel), and bar plot of percentage of significantly lengthened or shortened genes in each enriched gene set, GeneRatio is the proportion of differentially polyadenylated genes that belong to a known gene set. C. Coverage by RNA-seq reads of 3’UTRs of genes *cyb-1/2.2, cdk-1* and *par-5,* red lines mark the estimated proximal polyadenylation site.

### Differential Splicing play roles in germline development

We further performed differential splicing analysis using rMATs (Shen et al., 2014) to look for differential transcription of alternatively spliced isoforms of genes between neighboring stages along the oocyte developmental axis. Since rMATs could not account for batch effects, we performed the analysis with samples from Batch A (Figure 2A). We identified varying numbers of genes exhibiting significant splicing signals defined by rMATs, i.e. alternative 3’ splice site (A3SS), alternative 5’ splice site (A5SS), intron retention (IR), mixed exon usage (MXE), skipped exon usage (SE), between neighboring stages. Most notably, the S1 vs S2 and the S4 vs F3 comparisons yielded the most differential splicing usage with 58 genes and 54 genes exhibiting differential splicing, respectively (Figure 7A). Genes with differential splicing usages between the S1 vs S2 comparison are enriched for GO terms related to mitosis (Figure 7B), which is expected, given the fact that S1 contains the TZ regions (Figure 1C). However, other neighboring stages comparisons yielded no significantly enriched GO Biological Process terms. A few interesting examples are detailed as follows.

**Figure 7.**
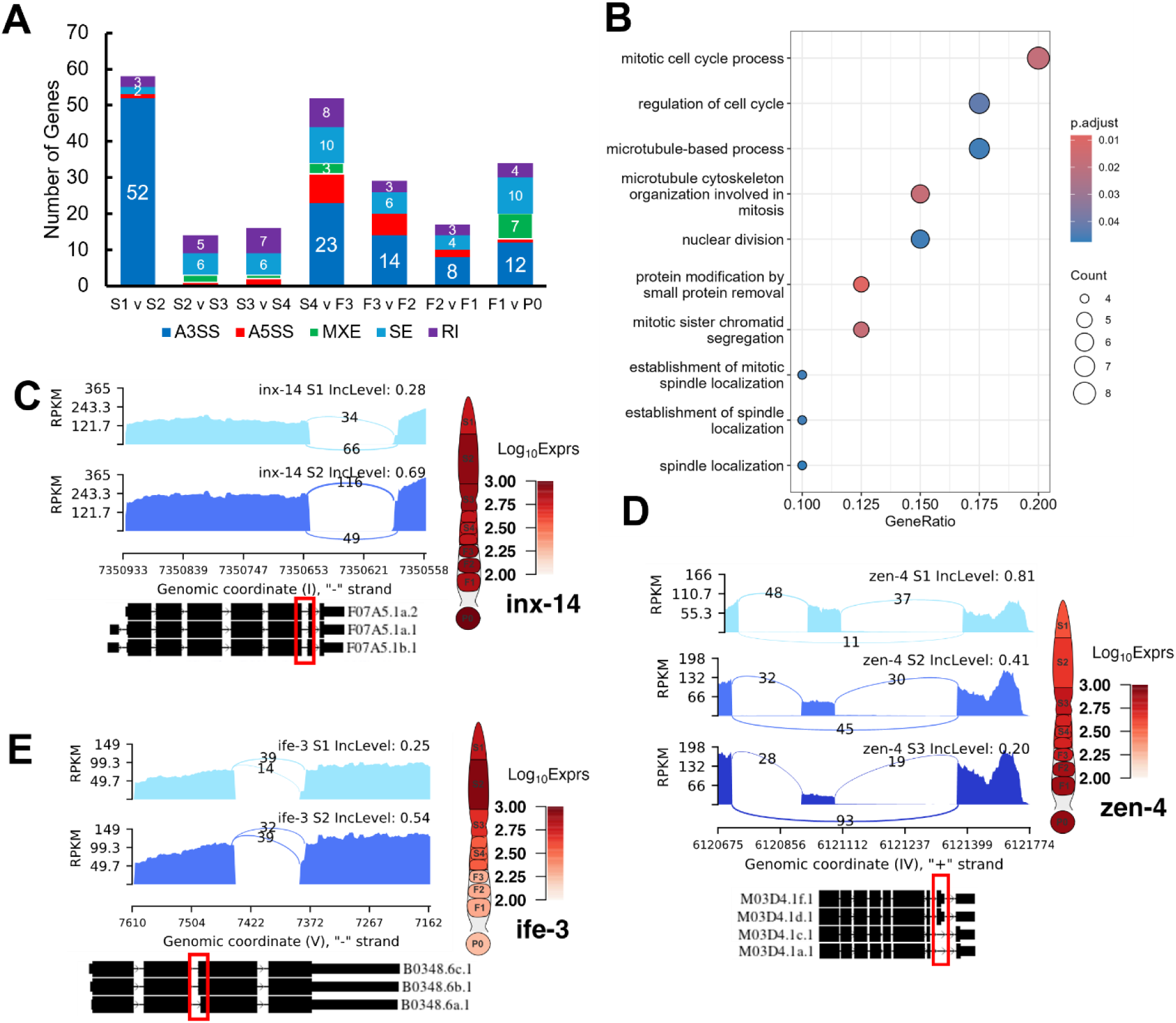
Examples of differential splicing usage of genes during germline development. A. Box plot of numbers of five splicing types (A3SS, A5SS, MXE, Se and RI, see main text for definitions) detected between each pair of neighboring stages of germline development. B. Enriched GO terms of genes with differential splicing events between the S1 and S2 stages, GeneRatio is the proportion of differentially spliced genes that belong to a known gene set. C. Differential splicing events of gene *inx*-14 between the S1 and S2 stages. D. Differential splicing events of gene *zen*-4 between the S1 and S2 as well as S2 and S3 stages. E. Differential splicing event of gene *ife*-3 between the S1 and S2 stages. Exact positions of splicing events are shown in the red box.

Gene *inx-*14 was differentially spliced during the S1 to S2 transition (Figure 7C). Specifically, *inx-*14 was preferentially utilized for its longer 6^th^ exon in the S2 stage compared to the S1 stage, resulting in increased proportions of its F07A5.1b isoform (Figure 7C). It has been documented that INX-14, along with INX-21/22 and INX-8/9 forms hemichannels that facilitate the communication between the somatic gonad and the germline and plays a role in meiosis to mitosis transitions by negatively regulating meiotic maturation and promoting germline proliferation (Starich et al., 2014). UniProt designates the F07A5.1b isoform as the canonical isoform, differing from the alternative F07A5.1a isoform by 2 amino acids in the 406-407 positions. Our results present a possible mechanism by which INX-14 changes its association with either INX-21 or INX-22 (Starich et al., 2014).

Another notable event was a gradual increase in preference of *zen-4* skipping its 8^th^ exon in the S1 to S3 transition (Figure 7D). Specifically, the most abundant *zen*-4 isoforms were M03D4.1a.1, M03D4.1c.1, M03D4.1d.1 and M03D4.1f.1 (Figure 7D). This is due to the lack of read coverage for the regions that are spanned by the other isoforms (Figure 7D). The exon skipping event is indicative of decreased preference for the M03D4.1d.1 and M03D4.1f.1 isoforms, which contain the skipped exon in the other isoforms (Figure 7D). ZEN-4 along with CYK-4 forms the centralspindlin complex, a conserved component of intercellular bridges that functions in the cellularization of cells during cytokinesis(Lee et al., 2018, Zhou et al., 2013, White and Glotzer, 2012). Though a recent study suggested that ZEN-4 was not essential in the germline for the closure of the intercellular bridge(Lee et al., 2018), our results suggest that as the oocyte moves along the rachis into late pachytene stage, alternative isoforms of *zen*-4 may still play a role in the cellularization of maturing oocytes.

In addition, we found that *ife*-3 switched isoforms during the S1 to S2 transition (Figure 7E). The *ife*-3 gene encodes one of the *C. elegans* homologs for human translation initiation factors (eIFs) that plays critical role in regulating mRNA content along with microRNA and RNA binding proteins(Huggins et al., 2020). More specifically, *ife*-3 functions as a repressor of *fem*-3 expression to promote production of oocytes in the germline(Huggins et al., 2020, Mangio et al., 2015). Here, we showed a switch in *ife*-3 splicing preference for the B0348.6b and B0348.6c isoforms over the shorter B0348.6a isoform (Figure 7E). Along with a slight increase in *ife*-3 expression, these results hint at a possible mechanism of *ife*-3 regulation in the pachytene stage of oogenesis. Interestingly, *ife*-3 expression reduced significantly in proximal oocytes, where transcription became increasingly silent, thus obviating the need for mRNA regulation (Figure 7E).

Other genes worth pointing out include *tos-1* coding for a reporter of differential splicing (Ma et al., 2011), and lev-11 coding for tropomyosin (Watabe et al., 2018). *Tos*-1 experiences loss of preference for the usage of its 3^rd^ exon from F1 to zygote (Supplementary Figure 4A), which is further corroborated by the decreased coverage of its longer isoform in S4 (Supplementary Figure 4A). However, the differential splicing of *lev*-11 transcripts (Supplementary Figure 4B) might occur in sheath cells wrapped around the oocytes as we argued earlier. It has been shown that different isoforms of *lev*-11 exhibit different characteristics in terms of muscle assembly and function(Watabe et al., 2018). Since F2 oocytes are roughly covered by Sh4 and F1 oocytes by Sh5, it is likely that an alternative isoform switch of *lev-*11 contributes to the different functions of these two sheath cell types.

## Discussion

With a spatial layout of cells that simultaneously mirrors the timeline of oogenesis, the *C. elegans* gonad can serve as a powerful model for uncovering mechanisms of oogenesis. However, the tiny size of the gonad also presents challenges for in-depth studies of the intricacies of this process. With the recent development of single cell methods, we utilize scRNA-seq techniques to decipher the transcriptomic landscape of different stages of oocyte formation as well as fertilization. Our transcriptomic dataset of the *C. elegans* gonad presents a good platform for research into the transcriptional landscape of oogenesis of animals. Our results not only are able to recall most of the oogenic genes designated by earlier research that utilized micro-arrays and bulk-RNAseq (Reinke et al., 2004, Ortiz et al., 2014), but also are highly correlated, through careful alignment of samples, with expression profiles of the different stages of the germline found by more recent studies that relied on single cell based techniques (Diag et al., 2018, Tzur et al., 2018).

Our expression profiles show a distinct pattern in the UMAP display, which is consistent with the developmental axis of the gonad, indicative of our successful capture of the transcriptomes underlying the oogenesis program. Though our dataset presents discernable batch effects, we either incorporated them into our analysis models or forfeited the smaller batch of samples when necessary. The number of biological repeats for each stage as well as the sequencing depth for each sample means the results are robust to the discarding of few samples.

Despite meticulous filtering of possible contamination of somatic tissues, the dissection of the tiny gonad presents a delicate problem, and it is difficult to fully avoid contamination by surrounding tissues. In this study, we note that distal stages (S1-S3) inevitably contain transcripts originating from Sh1 and Sh2 sheath cells, due to the unenclosed and miniscule nature of the germline along the rachis. Thus, we focus on elucidating the transcriptomic changes of known germline associated genes to minimize false positive findings. Despite separating proximal stage oocytes as best as possible from shattered somatic tissue, e.g., sheath cells. complete removal of somatic components remained an elusive task. This prompted us to investigate patterns of expression that may arise from known somatic specific genes and take care in interpreting the results. We find that a great portion of genes that are drastically downregulated in zygotes relative to the F1 oocyte are of somatic origin, including many known markers of muscle cells and sheath cells. This allowed us to mark a large portion of genes as somatic in nature, especially in the proximal gonad samples. However, we argue that the expression profiles of these genes depleted in zygotes are not without merit. From their expression patterns throughout the gonad, these genes can be divided into roughly two groups. The first consisting of genes that have relatively stable transcription in the proximal oocytes before complete disappearance in zygotes, and a second group consisting of genes that are drastically upregulated in only the F1 oocyte. This second group of genes includes those whose transcripts have recently been found to be produced in the spermathecae but transported into F1 oocytes (Trimmer et al., 2023). We thus provide a list of genes that might undergo this process. Though the exact function and underlying mechanisms for this phenomenon remain to be elucidated, a few genes exhibiting this pattern have been shown to affect the ageing of *C. elegans* (Zimmerman et al., 2015).

We confirm previous findings(Lee et al., 2012) at the transcriptomic level that the growth of oocytes presents as a process in which ribosomal biogenesis and cellular activity gradually decreases. Moreover, we observed at the transcriptional level known dynamics of core regulators of the mitosis to meiosis switch and meiosis maturation. In addition, we find that many genes involved in the eggshell formation initiate transcription as early as in the S1 stage, and their transcripts are accumulated until post fertilization. The *C. elegans* germline also presents a remarkable model for studying germ cell apoptosis(Gumienny et al., 1999)). Our results not only capture distinct upregulation of apoptosis related genes in the pachytene loop (S2 stage), but also discover novel candidate genes for future studies of germ line apoptosis. Furthermore, our gene clustering and DEG results also reveal three distinct sets of apoptotic related genes, characterized by the expression pattern of ced-3/4/9 (Figure 6C), *ced*-8 (Figure 6D) and *egl*-1(Figure 6E), respectively. These different modes of transcription suggest that cross-talks occur between different genes at the transcriptional and post-transcriptional levels to induce apoptosis.

The previous report that RBP and the 3’UTRs are key players in a complex regulatory mechanism (Merritt et al., 2008, Mangone et al., 2010b, Diag et al., 2018) in the *C. elegans* germline prompted us to investigate whether significant changes in polyadenylation site usage occurred during oogenesis and fertilization. Though we were not able to find significantly enriched pathways regulated via changes in polyadenylation during oogenesis, we did find enrichment for cell cycle processes due to changes in polyadenylation site usage during fertilization. Our results suggest that polyadenylation sites of transcripts remain relatively stable during oogenesis, and active regulation of alternative polyadenylation likely resumes in the zygote.

Finally, we find that alternative splicing events are present throughout the gonadal segments. We reveal significant changes in the usage of isoforms of hemi-channel gene *inx*-14. It is highly likely that products of different isoforms of *inx*-14 associate with germline hemichannels INX-21/22 or somatic hemichannels INX-8/9 to facilitate communication between the somatic gonad and germline. We also find differential splicing usage of genes in the germline. For instance, we observe differential splicing of *zen-*4 throughout the pachytene region. Though previous studies preclude the involvement of ZEN-4 in oocyte cellularization in the germline syncytium (Lee et al., 2018), ZEN-4 isoforms may still serve functions in oocyte growth in late pachytene. Future studies are needed to elucidate the roles of isoform usages in *C. elegans* oogenesis and the underlying mechanisms.

Taken together, our results paint a complex transcriptional landscape of the germline development, oogenesis and fertilization processes in *C. elegans* in finer detail than previous studies. Though contamination of somatic tissue presented challenges, we are able to discern putative somatic elements. We not only confirm previous findings, but also present many novel discoveries of transcriptional events along the temporospatial axis of the *C. elegans* germline and the zygote. Though much work remains to be done, particularly, with better dissection techniques to remove somatic contaminations, our study still presents a wealth of resources and gene candidates for future experimental investigation to reveal the underlying mechanisms of the oogenesis program.

## Materials and Methods

### Experimental Model

The AZ212 *C. elegans* strain was obtained from the *C. elegans* Genetics Center (University of Minnesota), and was maintained in E. coli OP50 lawn on an agar plate according to the standard protocol(Stiernagle, 2006).

### Method Details

#### Dissection of the gonad and harvest of samples

After a well-fed gravid hermaphrodite was immobilized in the egg salt solution (ESS) with 10% tetramisole (Sigma, St. Louis), a cut was made across the vulva using a 26G subcutaneous needle controlled by a micromanipulator (ROE-200, Sutter) under an inverted microscope (Olympus 1X71). This would release fertilized eggs and early-stage embryos from the uterus as well as sperms and at least portions of the two sides of the gonad. Each end of the gonad wrapped around by five pairs of sheath cells was completely isolated by pushing its distal end as shown in Figure 1A. The distal proliferative zone (S1) was cut off around the transition zone (Figure 1A) and harvested in about 10 nl ESS by suction using a patch clamp pipette under controlled of another micromanipulator (ROE-200, Sutter). The pachytene zone (S2), the loop corresponding to the diplotene zone (S3) and diakinesis zone (S4) were sequentially cut off at the positions as shown in Figure 1A and similarly harvested. The -3 (F3), -2 (F2), and -1 (F1) oocytes were also isolated by cutting through their boundaries and similarly harvested (Figure 1A). The zygote (fertilized oocyte) also known as P0 was similarly harvested when the two pronuclei were fused at its center (Figure 1A). Unavoidably, sheath cells wrapped around the gonad segments and oocytes as well as released sperms could be harvested in the samples.

### Preparation of RNA-seq libraries

We prepare a RNA-seq library for each harvested sample for Illumina platforms using a modified scRNA-seq method based on Tang et al as previously described (Su et al., 2023, Tang et al., 2010a, Tang et al., 2010b) at the earlier stage of the project and using the Smart-seq2 protocol (Picelli et al., 2013) later on. The libraries were sequenced by 100 bp paired-end reads on an Illumina HiSeq2000 or HiSeq2500 machine.

### Transcriptome Mapping and Quantification

The *C. elegans* genome assembly (GCA_000002985.3) was obtained from NCBI Refseq, while the annotations were based on Wormbase version: WS291. Prior to mapping, raw reads were trimmed with Trim Galore(Krueger, 2015), with parameters (quality >= 10, length > 35bp). We quantified the expression levels of genes in two ways for different subsequent analysis. For differential gene expression analysis, trimmed reads were mapped to the genome using HISAT2(Kim et al., 2019) with default settings, read counts were obtained by using HTSeq (Anders et al., 2015) with default settings based on the mapping results. The trimmed reads were also mapped to the genome using Salmon (Patro et al., 2017) with default settings to obtain transcript per million (TPM) estimates for both genes and transcripts.

### Quality Control

Sequenced libraries were then assessed for quality with custom scripts and quality metrics evaluated via the QoRTs package(Hartley and Mullikin, 2015). First, Salmon quantified TPM values for mitochondrial genes, spike-ins and sperm specific genes (See also, Supplementary Figure S1, Supplementary Table S2) were obtained. A sample was filtered out if it met any of the following criteria: i) over 5% reads (in terms of TPM) were from the mitochondrial genome; ii) over 5% reads (in terms of TPM) were from rRNA genes; iii) over 5% reads (in terms of TPM) were from sperm specific genes); iv) over 5% reads (in terms of TPM) were from intestine specific genes; v) HISAT unique reads mapping rate < 70%; vi) less than 50% of HISAT uniquely mapped reads were mapped to coding DNA sequences (detailed procedure and genes that were removed and used for filtering are shown in Supplementary figure S1 and Supplementary Table S2). These criteria were set to remove samples that were of poor libraries quality or were heavily contaminated by sperm, intestinal tissue and/or exhibited reduced quality during sample collection. To further increase the robustness of subsequent analysis, samples were visualized using Uniform Manifold Approximation and Projection (UMAP), and those that largely deviated from clustered groups of the same sample type were removed. In addition, we included the P0 (1-cell) samples of Tintori, et al (Tintori et al., 2016) in our analysis, and the samples were processed through the same pipeline as our own samples.

### Comparison with Previous Datasets

Gene expression data from six previous studies were collected from the following sources and compared with our data. For all comparisons, we used filtered samples with all genes (genes were not filtered). Details of the datasets and comparisons are as follows:

1. Reinke et al. 2004 (Reinke et al., 2004) provided the first microarray-based list of oogenic genes. The list was retrieved from via their supplementary material.
2. Ortiz et al. 2014 (Ortiz et al., 2014) performed RNA-seq analysis on the gonad to distill a list of genes termed oogenic. These genes were acquired via their supplementary data, and genes marked oogenic were used for our subsequent comparisons.
3. Stoeckius et al. 2014 (Stoeckius et al., 2014) performed RNA-seq on proximal oocytes and 1 cell zygotes. Expression profiles were acquired via the instructions in their paper and genes with expression > 0.5 RPKM were deemed expressed.
4. West et al, 2018 (West et al., 2018) dissected the gonad into mitotic and meiotic sections, and oocytes. RNA-seq data of each sample was acquired via the supplementary material of the paper, and genes with a reads count > 0 were deemed expressed.
5. Tzur et al. 2018 (Tzur et al., 2018) utilized the Cel-seq protocol to sequence 10 segments of the *C. elegans* gonad, with 2 replicates per segment. Alignment of these 10 segments to our segments was based on diagrams presented in their study and rough estimates of where their dissection occurred. The exact alignments between their segments and ours are given in Supplementary Table S1. Count matrices were acquired per the authors’ instructions. Pearson correlation was performed with log transformed count values using all shared genes.
6. Diag et al, 2018 (Diag et al., 2018) performed cryo-dissection of the 3 posterior and 3 anterior gonads into 13-15 segments per gonad. This resulted in 85 slices sequenced via Cel-seq. Expression profiles for these samples were retrieved from GEO with accession number GSE115884. Samples with < 10^4^ reads were discarded from correlation analysis with our samples. The authors (Diag et al., 2018) provided approximate slice label, slice size as well estimates size of each gonad region. Thus, we were able to derive a coarse conversion from their slices to our segments, as shown in Supplementary Table S1. Pearson correlation was performed with log transformed count values using all shared genes.

### Differential Gene Expression Analysis

We performed differential gene expression analysis between each two dissected neighboring stages along the developmental axis of the gonads as above-described and zygotes using Monocle2 (Qiu et al., 2017). Experimental batch and gene detection rate in each sample were included as covariates along with segment/cell-type to model normalized gene expression using the negbinomial.size model of Monocle2. Because Monocle2 does not produce Log2FoldChange values, we applied Bayesian shrinkage of gene model coefficients using the apeglm (Zhu et al., 2019) package to account for large foldchange values of genes with low expression and obtain shrunken Log_2_FC values for each gene. A model of gene expression as a function of segment/cell-type was also fit to assess genes that were differentially expressed across all stages prior to fertilization (excluding P0). Genes with an FDR < 0.05 and a fold change increase/decrease of 1.5 were considered differentially expressed. ClusterProfiler(Yu et al., 2012) was used to perform Gene Set Enrichment Analysis (GSEA) with pre-ranked shrunken Log2FC values and gene sets from KEGG(Kanehisa et al., 2022), GO(Gene Ontology, 2021) Biological Pathways, Reactome(Milacic et al., 2024) and Wikipathways(Martens et al., 2021). Enrichment of each type of gene sets was performed separately, and the results were aggregated. Only gene sets containing more than 10 and less than 250 genes were considered, and those with an FDR < 0.05 were considered significantly enriched.

### Clustering co-expressed genes

The union of DEGs identified in all pairwise comparisons were used for gene co-expression analysis. After the read count values of genes were variance stabilizing transformed using the vstExprs function of Monocle2 package, Pearson correlation coefficient between expression levels of the genes in the samples were calculated, and genes were hierarchically clustered using the ‘ward.D2’ method of the hclust function in R. Upon visual inspection of the resulting clustering heatmaps, the clusters were set at a hierarchical level. Each cluster was then subject to enrichment analysis for GO biological process (BP) terms using ClusterProfiler (Yu et al., 2012) to identify significantly enriched terms for the cluster. Gene expression as well as the respective clusters were visualized with the ComplexHeatmap package(Gu, 2022), and the top three most significantly (fdr < 0.05 or p-value < 0.001) enriched GO terms were shown alongside the heatmap.

### Differential Alternative Polyadenylation Analysis

3’UTR regions were extracted from the WS291 annotation via custom scripts to only include 3’UTR regions that did not overlap coding exons and other UTR regions. The samtools (Li et al., 2009) depth function was used to obtain pair-read aware coverage of the genome for each samples with HISAT2(Kim et al., 2019) aligned bam files. Coverage for each sample was normalized with DESeq2(Love et al., 2014) size factors before estimation of polyadenylation site and long/short 3’UTR coverage, and Percentage of Distal polyA site Usage Index (PDUI) was computed performed using DaPars2(Feng et al., 2018, Li et al., 2021). Modification to the DaPars2 program was made to begin polyadenylation site search starting from 25bp downstream a 3’UTR’s 5’ end. For each neighboring stages comparison, only 3’UTRs that belonged to a gene with a mean count >10 across all compared samples and had PDUI values in at least 3 samples in both stages were tested for differential alternative polyadenylation. Fisher’s exact test was performed with the average long/short 3’UTR coverage in compared stages, and the resulting p-values were corrected for false discovery rate (FDR) via the Benjamini Hochberg method. Genes that had FDR < 0.05 and |PDUI difference| > 0.05 were called for significantly differential alternative polyadenylation. ClusterProfiler(Yu et al., 2012) was used to perform GO BP (Ashburner et al., 2000) term enrichment analysis, and significant terms with FDR < 0.05 were called significantly enriched. Visualization was made with the trackViewer(Ou and Zhu, 2019) R package.

### Differential Splicing Analysis

Differential splicing analysis was performed using rMATS (Shen et al., 2014) that calculated splicing Psi values and evaluated their statistical significance. rMATS classifies splicing events into five categories: alternative 3’ splice site (A3SS), alternative 5’ splice site (A5SS), intron retention (IR), mixed exon usage (MXE), skipped exon usage (SE). A splicing event with a Psi value change > 0.1 and an adjusted p-value < 0.05 was considered to be significant.

### Resource availability

#### Lead Contact

Further information and requests for resources and reagents should be directed to and will be fulfilled by the lead contact, Zhengchang Su

#### Materials availability

No new reagents were generated from this paper.

#### Availability of data and materials

- Single-cell RNA-seq data have been deposited at GEO Accession: GSE261784 and are publicly available as of the date of publication.
- All original code has been deposited at https://github.com/bio-info-guy/Celegans_paper/ and is publicly available as of the date of publication.
- Any additional information required to reanalyze the data reported in this paper is available from the lead contact upon request.

## Competing Interest Statement

The authors declare that they have no competing interests.

## Acknowledgements

The work was supported by the US National Science Foundation (DBI-1661332) and NIH (R01GM106013). The funding bodies played no role in the design of the study and collection, analysis, and interpretation of data and in writing the manuscript. We would also like to thank Willian Su and Richard Cai for help in the earlier stage of the project for data generation and all lab members for their discussion and suggestions.

## Authors’ contributions

ZS conceived the project. JS and DD generated the scRNA-seq data, YS performed data analyses. YS and ZS wrote the manuscripts. All authors read and approved the final manuscript.

## Notes

### Competing Interest Statement

The authors have declared no competing interest.

