## Supplementary Information for "Transcriptomic Analysis of the Spatiotemporal Axis of Oogenesis and Fertilization in *C. elegans*"

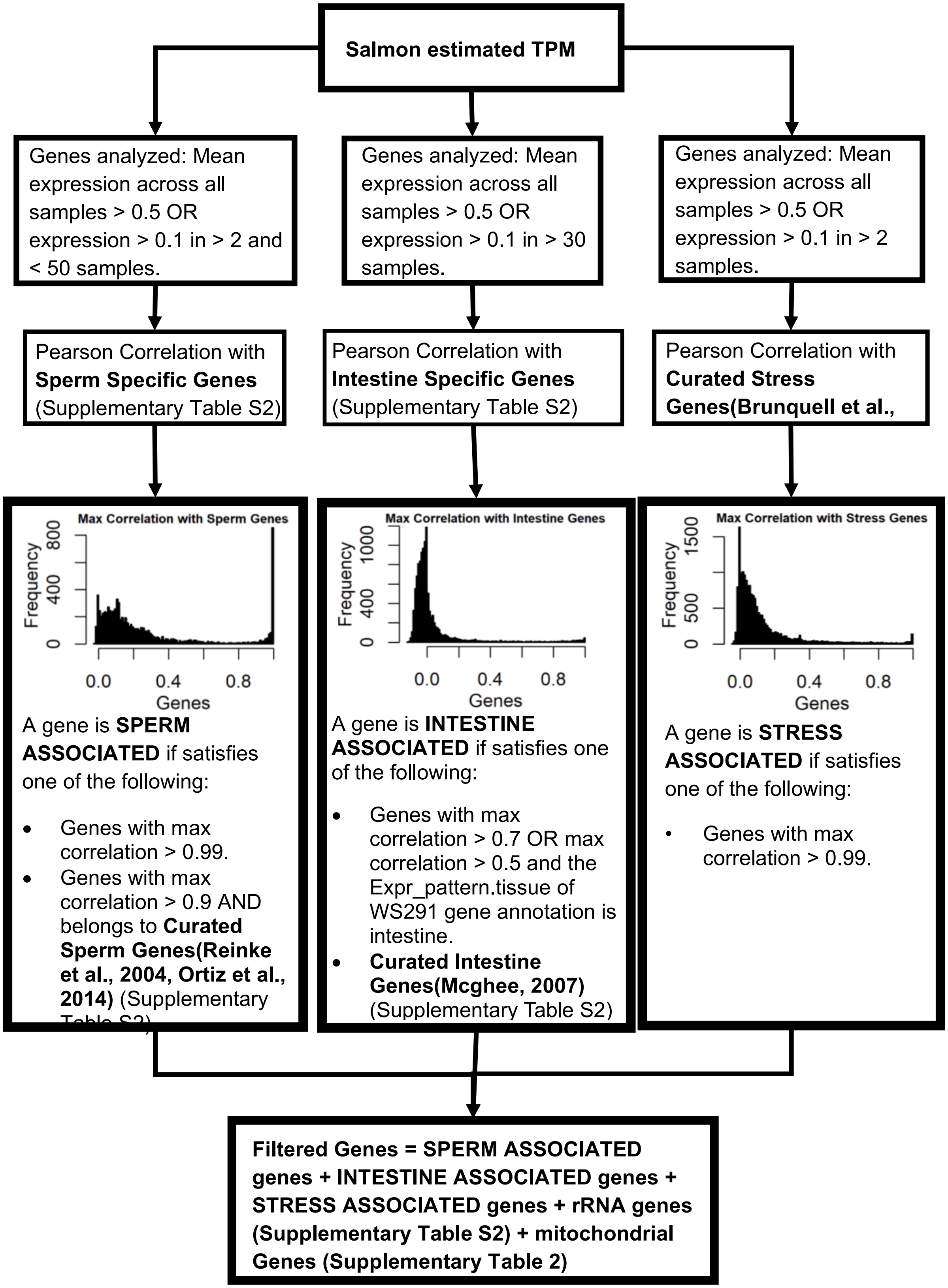

Salmon estimated TPM
Genes analyzed: Mean expression across all samples > 0.5 OR expression > 0.1 in > 2 and < 50 samples.
Genes analyzed: Mean expression across all samples > 0.5 OR expression > 0.1 in > 30 samples.
Genes analyzed: Mean expression across all samples > 0.5 OR expression > 0.1 in > 2 samples.
Pearson Correlation with Sperm Specific Genes (Supplementary Table S2)
Pearson Correlation with Intestine Specific Genes (Supplementary Table S2)
Pearson Correlation with Curated Stress Genes(Brunquell et al.,
Max Correlation with Sperm Genes
Frequency
0
400
800
0.0
0.4
0.8
Genes
Max Correlation with Intestine Genes
Frequency
0
400
1000
0.0
0.4
0.8
Genes
Max Correlation with Stress Genes
Frequency
0
500
1500
0.0
0.4
0.8
Genes
A gene is SPERM ASSOCIATED if satisfies one of the following:
• Genes with max correlation > 0.99.
• Genes with max correlation > 0.9 AND belongs to Curated Sperm Genes(Reinke et al., 2004, Ortiz et al., 2014) (Supplementary Table S2)
A gene is INTESTINE ASSOCIATED if satisfies one of the following:
• Genes with max correlation > 0.7 OR max correlation > 0.5 and the Expr_pattern.tissue of WS291 gene annotation is intestine.
• Curated Intestine Genes(Mcghee, 2007) (Supplementary Table S2)
A gene is STRESS ASSOCIATED if satisfies one of the following:
• Genes with max correlation > 0.99.
Filtered Genes = SPERM ASSOCIATED genes + INTESTINE ASSOCIATED genes + STRESS ASSOCIATED genes + rRNA genes (Supplementary Table S2) + mitochondrial Genes (Supplementary Table 2)

**Supplementary Figure S1 Gene filtering pipeline for mitochondrial, rRNA, intestine, sperm and stress associated genes, related to Methods and Materials**. WS291 gene annotations are obtained via the Wormbase SimpleMine tool(Harris et al., 2019, Davis et al., 2022).

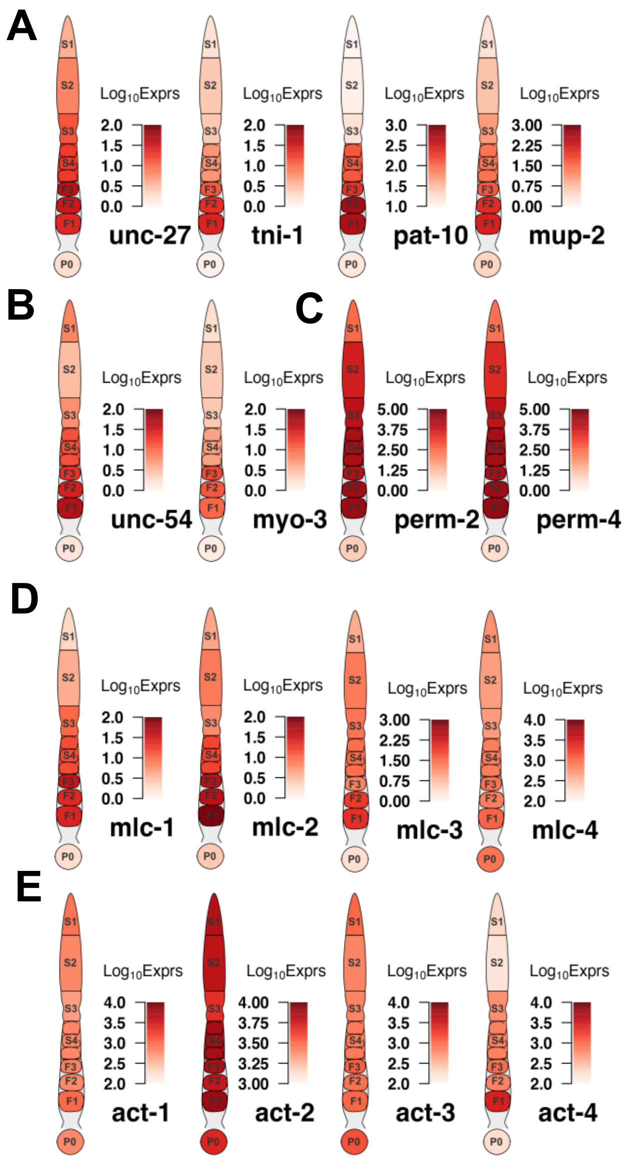


**Supplementary Figure S2.** A. genes encoding for components of the troponin complex. B. Genes coding for myosin heavy chain proteins (unc-54 and myo-3). C. Genes coding eggshell components perm-2/4. D. Genes coding for four myosin light chains encoding genes. E. Genes coding for four actin encoding genes.

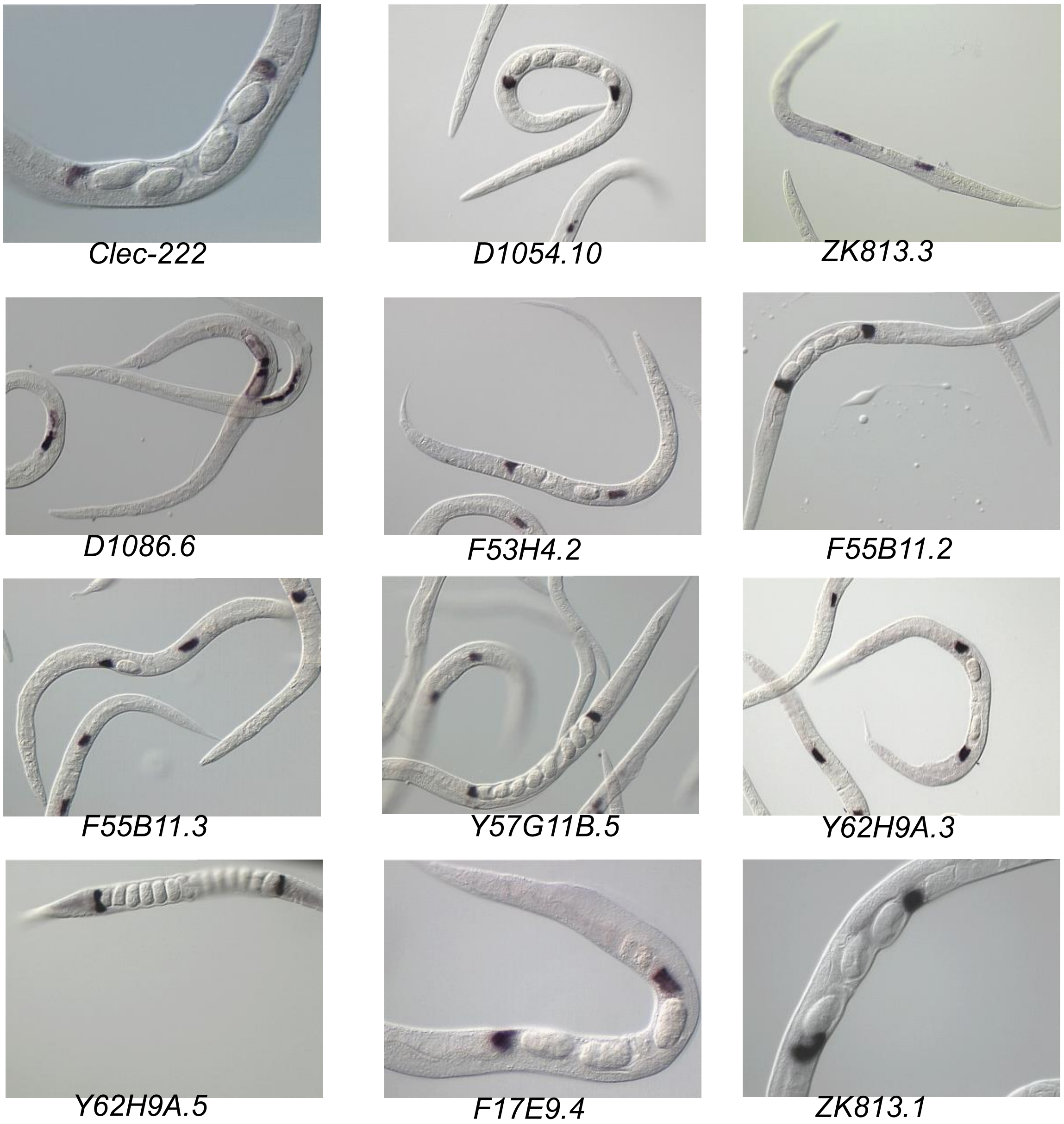


**Supplementary Figure S3. NEXTDB(Kohara, 2001) in situ imaging of 13 of the 25 putative genes that originate from the spermathecae, related to Figure 5C and Results .**

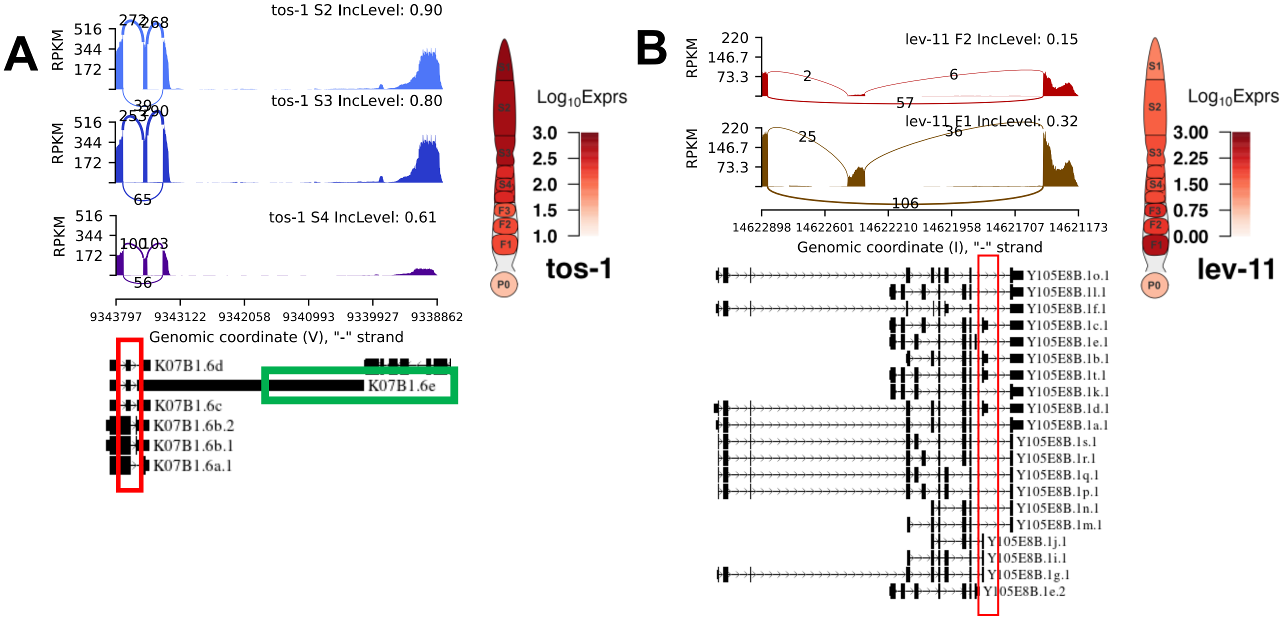


**Supplementary Figure S4.** A. Differential splicing events of tos-1 between S2 and S3 as well as S3 and S4 stages. B. Differential splicing event of lev-11 between F2 and F1 oocytes. Exact positions of splicing events are shown in the red box, long isoform of tos-1 is shown in green box.

| Our segments | Tzur et al, 2018(Tzur et al., 2018) | Diag et al, 2018(Diag et al., 2018) |
|---|---|---|
| **S1** | Segments 1,2 | Segments 1,2 |
| **S2** | Segments 3,4,5 | Segments 4, 5, 6, 7 |
| **S3** | Segments 6,7 | Segment 8 |
| **S4** | Segment 8 | Segments 10,11 |
| **F3** | Segment 9 | Segment 12 |
| **F2** | Segment 9 | Segment 13 |
| **F1** | Segment 10 | Segments 14, 15 |
| **P0** | NA | NA |

**Supplementary Table S1. Correspondence between or gonad stages and that of Diag et al, 2018(Diag et al., 2018) and Tzur et al, 2018(Tzur et al., 2018), related to Methods and Materials.**

**Supplementary Table S2. Collection of mitochondrial, sperm, intestine, stress and rRNA genes used for filtering, related to Methods and Materials.** Curated Sperm Genes are obtained from (Reinke et al., 2004, Ortiz et al., 2014), Curated Intestine genes are obtained from (Mcghee, 2007), Curated stress genes are obtained from (Brunquell et al., 2016).

**Supplementary Table S3. Differential expression results for pairwise comparison between neighboring stages, related to Figure 3C.** Stages.Tested column indicates which 2 stages are compared.

**Supplementary Table S4. Gene cluster membership for all significant DEGs, related to Figure 4.**

**Supplementary Table S5**. **Significantly enriched gene sets from GSEA analysis on differential expression results of each comparison, related to Results.** Stages.Tested column indicates which 2 stages are compared.

| **Putative Genes from Spermathecae** |
| --- |
| *clec-222, D1054.10, ule-3, D1086.6, F53H4.2, F54F7.3, F55B11.2, F55B11.3, F57C2.4, K07A1.6, Y37D8A.19, Y57G11B.5, Y62H9A.3, Y62H9A.4, Y62H9A.5, ule-5, ZC373.2, E02H9.7, F17E9.4, ZK813.1, ZK813.3, D1086.11, H29C22.1, ZK813.7, F38A5.22* |

**Supplementary Table S6**. **25 putative genes that originate from the spermathecae, related to Figure 5C and Results.** Genes with high expression in F1 cells that are significantly upregulated between F2 and F1 and significantly downregulated between F1 and P0.
